# Arabidopsis PAD4 lipase-like domain is a minimal functional unit in resistance to green peach aphid

**DOI:** 10.1101/769125

**Authors:** Joram A. Dongus, Deepak D. Bhandari, Monika Patel, Lani Archer, Lucas Dijkgraaf, Laurent Deslandes, Jyoti Shah, Jane E. Parker

## Abstract

Plants have evolved mechanisms to attract beneficial microbes and insects while protecting themselves against pathogenic microbes and pests. In Arabidopsis, the immune regulator PAD4 functions with its cognate partner EDS1 to limit pathogen growth. PAD4, independently of EDS1, reduces infestation by Green Peach Aphid (GPA). How PAD4 regulates these defense outputs is unclear. By expressing the N-terminal PAD4-lipase-like domain (LLD) without its C-terminal ‘EDS1-PAD4’ (EP) domain, we interrogated PAD4 functions in plant defense. Here we show that transgenic expression of PAD4^LLD^ in Arabidopsis is sufficient for limiting GPA infestation, but not for conferring basal and effector-triggered pathogen immunity. This suggests that the C-terminal PAD4-EP domain is necessary for EDS1-dependent immune functions. Moreover, PAD4^LLD^ is not sufficient to interact with EDS1, indicating the PAD4-EP domain is required for heterodimerisation. These data provide molecular evidence that PAD4 has domain specific functions.

## Introduction

To colonize plants, pathogenic microbes and pests (such as aphids or nematodes) deliver susceptibility factors, called effectors, to the host which target defenses and reprogram cells to promote infection or infestation. Many host-adapted biotrophic and hemi-biotrophic pathogens deploy effectors to disable PAMP/MAMP-triggered immunity (PTI) mediated by cell surface-resident receptors [Boutrot & Zipfel, 2017; Dangl & Jones, 2006; Dodds & Rathjen, 2010]. These microbes encounter two further important immunity barriers. One is conferred by intracellular nucleotide-binding leucine-rich repeat (NLR) receptors recognizing interference by specific effectors [Jones et al., 2016]. NLR activation leads to effector-triggered immunity (ETI) involving the rapid transcriptional mobilization of resistance pathways and, often, localized host cell death, which limit pathogen infection [Bhandari et al., 2019; Cui et al., 2015; Mine et al., 2017]. NLR-mediated immune responses are also effective against probing insects and nematodes [Milligan et al., 1998; Rossi et al., 1998; Villada et al., 2009; Wroblewski et al., 2007]. A second barrier, called basal immunity, slows virulent pathogen growth and disease progression by eliciting a weak immune response [Cui et al., 2015; Cui et al., 2017; Dangl & Jones, 2006]. Although the precise activation mechanism for post-infection basal immunity is not known, in Arabidopsis it requires several ETI signaling components [Century et al., 1995; Feys et al., 2001; Glazebrook et al., 1997; Parker et al., 1996], and is proposed to be the culmination of weak NLR-triggered ETI combined with residual PTI [Cui et al., 2017; Gantner et al., 2019].

In Arabidopsis, the nucleocytoplasmic immune regulator PAD4 (PHYTOALEXIN DEFICIENT4) signals in both ETI and basal immunity by stimulating production of the defense hormone salicylic acid (SA) and anti-microbial molecules, which limit pathogen growth [Glazebrook et al., 1997; Jirage et al., 1999; Wiermer et al., 2005; Zhou et al., 1998]. PAD4 is a member of a small family (the EDS1 family) of sequence-related immunity regulators, comprising also EDS1 (ENHANCED DISEASE SUSCEPTIBILITY1) and SAG101 (SENESCENCE ASSOCIATED GENE101) [Feys et al., 2005; Lapin et al., 2019]. Arabidopsis EDS1 and PAD4 function together in conferring ETI governed by a sub-class of NLRs with N-terminal Toll-interleukin1 Receptor domains (known as TIR-NLRs or TNLs) [Feys et al., 2005; Jones et al., 2016; Wagner et al., 2013]. Genetic and molecular studies in Arabidopsis revealed that activated TNL receptors stimulate EDS1–PAD4 basal immunity activity to transcriptionally boost SA signaling and other defense responses, and repress antagonistic jasmonic acid (JA) hormone pathways [Cui et al., 2017; Cui et al., 2018]. In Arabidopsis, the EDS1-PAD4 transcriptional reprogramming function in pathogen immunity requires a nuclear EDS1 pool [Bartsch et al., 2006; Cui et al., 2017; Garcia et al., 2009; Stuttmann et al., 2016].

EDS1, PAD4 and SAG101 each possess an N-terminal lipase-like domain (LLD) with an α/β hydrolase topology resembling eukaryotic class-3 lipase enzymes [Rauwerdink & Kazlauskas, 2015; Wagner et al., 2013; Wang et al., 2018], and a structurally unique C-terminal EP (EDS1-PAD4) domain consisting of α-helical bundles [PFAM database: PF18117; Wagner et al., 2013]. The EDS1 and PAD4, but not SAG101, LLDs have a canonical Ser-Asp-His (S-D-H) catalytic triad that is characteristic for α/β hydrolases [Wagner et al., 2013]. The serine is part of a characteristic lipase GXSXG motif which is conserved in EDS1 and PAD4 proteins across seed plant (angiosperm and gymnosperm) species [Wagner et al., 2013; Lapin et al., 2019]. Strikingly, the S-D-H residues were found to be dispensable for EDS1 and PAD4 signaling in Arabidopsis TNL-mediated ETI and basal immunity, indicative of a non-catalytic mechanism in pathogen resistance [Louis et al., 2012; Wagner et al., 2013].

EDS1 forms stable and mutually exclusive heterodimers with PAD4 or SAG101, consistent with distinct roles of these two EDS1 complexes in immunity [Lapin et al., 2019; Rietz et al., 2011; Wagner et al., 2013]. Based on a structural model of the EDS1-PAD4 heterodimer generated from the *At*EDS1-*At*SAG101 crystal structure, analysis showed that the juxtaposed LLDs are major drivers of heterodimerisation, likely promoting association of the aligned EP domains to form a cavity [Wagner et al., 2013]. The *At*EDS1^LLD^ alone, although stable, did not confer pathogen resistance, indicating that its EP domain is crucial for immune signaling activity [Wagner et al., 2013]. Further structure-based analysis identified an *At*EDS1 EP-domain surface lining the EDS1-PAD4 heterodimer cavity which is essential for the rapid transcriptional reprogramming of host cells in Arabidopsis TNL ETI [Bhandari et al., 2019; Lapin et al., 2019].

In Arabidopsis, *PAD4* mediates resistance to green peach aphid (GPA, *Myzus persicae* Sülzer) independently of *EDS1* and *SAG101* [Pegadaraju et al., 2005 & 2007]. GPA population growth was higher on Arabidopsis *pad4* compared to wild-type (WT) and *eds1, sag101* or *eds1/sag101* mutant plants [Pegadaraju et al., 2007]. Notably, *PAD4*-mediated defenses against GPA were found to not involve SA or camalexin production [Pegadaraju et al., 2005]. Moreover, in contrast to basal immunity and ETI, resistance to GPA was dependent on the S-D-H predicted catalytic triad residues PAD4^S118^ and PAD4^D178^, but not PAD4^H229^ [Louis et al., 2012; Wagner et al., 2013]. These different requirements suggest that PAD4 functions in immunity as a heterodimer with EDS1 are distinct from its function in resistance to GPA.

To gain a deeper insight into the molecular function of PAD4, we investigate here the properties of the PAD4^LLD^ in resistance to GPA and pathogen immunity. We show that the PAD4^LLD^ alone is sufficient to control GPA infestation, independently of EDS1 association. By contrast, we find that the Arabidopsis PAD4^LLD^ is insufficient for EDS1-dependent basal immunity and ETI, indicating that, like EDS1, the PAD4 EP domain is crucial for inducing immunity pathways. These results suggest that PAD4 can operate as a bipartite protein with the LLD and EP domains carrying out distinctive and separable roles in plant defense.

## Results

### The PAD4^LLD^ protein accumulates *in planta*, but does not interact with EDS1

*At*EDS1-*At*PAD4 heterodimer formation is driven chiefly by an N-terminal EDS1 hydrophobic loop (α-helix H; EDS1^LLIF^) and the juxtaposed PAD4^MLF^ motif (Figure 1A-C) [Feys et al., 2001; Wagner et al., 2013]. To test PAD4^LLD^ properties, we generated an *At*PAD4^LLD^ protein (residues 1-299; Figure 1A; blue). Transient overexpression of GFP-tagged PAD4^LLD^ in *Nicotiana benthamiana* produced a stable protein that did not co-immunoprecipitate (co-IP) FLAG-tagged EDS1, whereas full-length GFP-PAD4 did (Figure 1D). Similarly, PAD4^LLD^ failed to interact with EDS1 in an *N. benthamiana* split-luciferase assay (Figure S1). These data suggest that a stable interaction between PAD4 and EDS1 *in planta* requires part or all of the PAD4 EP domain in addition to the LLD interface.

**Figure 1.**
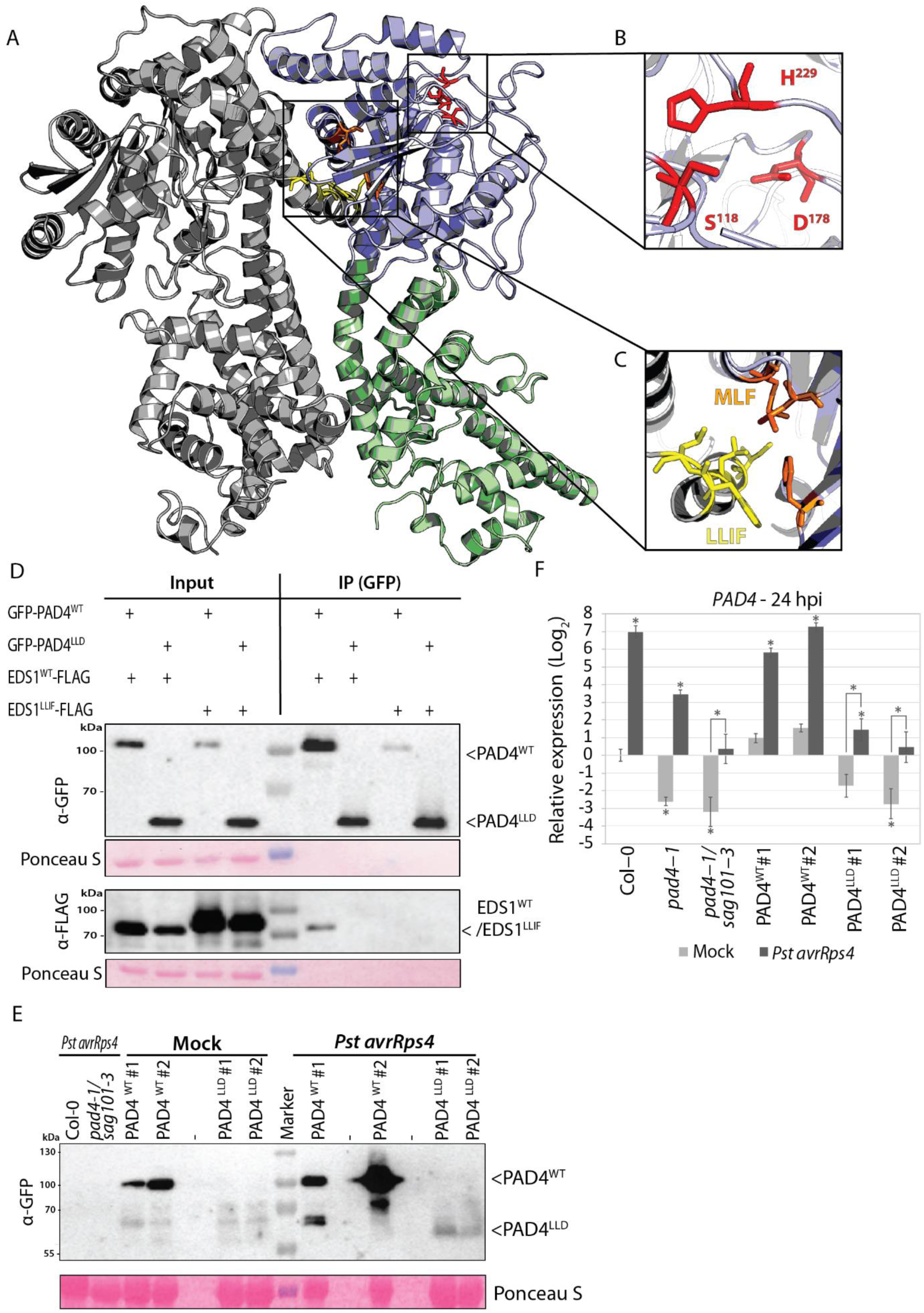
PAD4^LLD^ accumulates *in planta* but does not interact with EDS1. **A**. EDS1-PAD4 heterodimer model, based on the *At*EDS1-*At*SAG101 crystal structure [Wagner et al., 2013]. EDS1 (Grey), PAD4 (LLD) (blue) and PAD4 EP domain (green) are represented in cartoon format. **B**. PAD4 catalytic triad residues S^118^, D^178^ and H^229^ in the LLD are shown as red sticks. **C**. EDS1-PAD4-interacting motifs EDS1^LLIF^ and PAD4^MLF^ are colored with yellow and orange sticks, respectively. **D**. Co-immunoprecipitation (GFP-trap) of GFP-PAD4/PAD4^LLD^ with EDS1/EDS1^LLIF^-3xFLAG transiently expressed in *N. benthamiana* leaves (using *35S::GFP-PAD4/PAD4*^*LLD*^ and *35S::EDS1/EDS1*^*LLIF*^*-3xFLAG* constructs, respectively) A representative image from three independent experiments is shown. **E**. PAD4 accumulation in independent stable transgenic Arabidopsis lines expressing YFP-PAD4^WT^ and YFP-PAD4^LLD^ probed by Western blotting using α-GFP antibody at 24 hpi with mock (10 mM MgCl2) or *Pst* AvrRps4 treatments. A representative image from three independent experiments is shown. **F**. *PAD4* transcript abundance was determined by RT-qPCR at 24 hpi in mock- or *Pst* AvrRps4-treated samples of the indicated Arabidopsis lines. Data are pooled from three independent experiments each with two to three biological replicates (n = 6–9). Bars represent means of three experimental replicates ± SE. Relative expression and significance level is set to Col-0 mock-treated samples. Asterisk indicates *p* < 0.01, one-way ANOVA with multiple testing correction using Tukey-HSD.

To investigate PAD4^LLD^ properties in Arabidopsis, we introduced WT PAD4 (*pPAD4*::s*trepII-YFP-cPAD4*^*WT*^) or PAD4^LLD^ (*pPAD4*::s*trepII-YFP-cPAD4*^*LLD*^) constructs into a *pad4-1/sag101-3* mutant (Col-0 accession). PAD4^LLD^ in two independent stable transgenic lines showed a nucleocytoplasmic localization similar to PAD^WT^ at 24 h post infection (hpi) with *Pseudomonas syringae* pv. *tomato* strain DC3000 expressing the effector avrRps4 (*Pst avrRps4*) (Figure S2). Delivery of avrRps4 by *Pst* triggers ETI in Col-0 mediated by the receptor pair RRS1-S/RPS4 (RESISTANCE TO RALSTONIA SOLANACEARUM1-S/RESISTANCE TO PSEUDOMONAS SYRINGAE4) [Birker et al., 2009; Heidrich et al., 2011; Narusaka et al., 2009; Saucet et al., 2015]. The PAD4^LLD^ distribution is in line with previously described nucleocytoplasmic localizations of EDS1^LLD^ and PAD4^LLD^/SAG101^EP domain^ chimeras *in planta* [Lapin et al., 2019; Wagner et al., 2013]. PAD4^LLD^ protein was also immuno-detected from leaf samples treated with *Pst avrRps4*, although at much lower levels compared to PAD4^WT^ lines (Figure 1E). This contrasts with similar PAD4^LLD^ and PAD4^WT^ accumulation in *N. benthamiana* transient assays (Figure 1D). Lower PAD4^LLD^ protein accumulation than PAD4^WT^ in mock- and *Pst avrRps4*-treated Arabidopsis leaves can be attributed in part to lower accumulation of *PAD4* transcripts in the PAD4^LLD^ compared to PAD4^WT^ transgenic lines (Figure 1F). Hence, the LLD domain of PAD4 is sufficient to maintain a WT-like nucleocytoplasmic localization, but loss of the EP domain substantially reduces PAD4-EDS1 interaction and PAD4 steady-state levels in Arabidopsis.

### Expression of PAD4^LLD^ confers GPA resistance

PAD4 acts independently of EDS1 to restrict aphid infestation, and this function is dependent on the PAD4^LLD^ located S^118^ and D^178^ predicted α/β-hydrolase catalytic triad residues (Figure 1A & 1B) [Louis et al., 2012; Pegadaraju 2007]. Since PAD4^LLD^ accumulates in Arabidopsis, we tested whether the PAD4^LLD^ alone is able to resist GPA infestation. Consistent with earlier data [Louis et al., 2012; Pegadaraju et al., 2007], *pad4-1, pad4-1/sag101-3* and a PAD4^S118A^ line (in *pad4-1/eds1-2/EDS1*^*SDH*^;Wagner et al., 2013) permitted a significant increase in aphid population size compared to Col-0 in a no-choice bioassay, indicating compromised resistance to GPA infestation (Figure 2). The PAD4^LLD^ lines resisted GPA to similar levels as PAD4^WT^ and Col-0, even though they expressed very low PAD4^LLD^ amounts (Figure 2). Hence, low steady state accumulation of PAD4^LLD^ protein (Figure 1E) is sufficient to counter GPA infestation in Arabidopsis, implying that PAD4^LLD^ has an *in planta* activity. Based on these data we conclude that PAD4^LLD^ is a stable protein entity able to confer GPA resistance.

**Figure 2.**
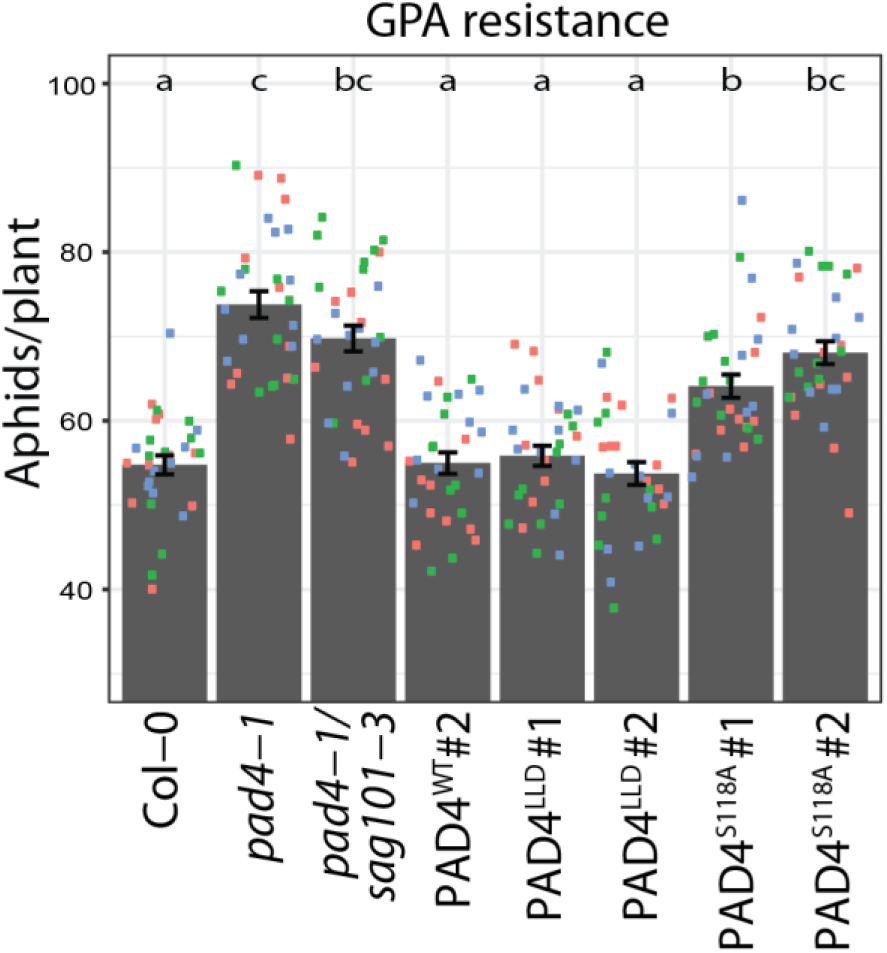
PAD4^LLD^ is sufficient for GPA resistance. Numbers of green peach aphids (GPA) per plant at 11 days post-infestation in a no-choice assay. Data are pooled from three independent experiments each with ten biological replicates per experiment (n = 30). Squares of the same color represent ten biological replicates in an independent experiment. Bars represent mean of three experimental replicates ± SE. Differences between genotypes were determined using ANOVA (Tukey-HSD, *p* <0.01), letters indicate significance class.

### Arabidopsis ETI and basal pathogen immunity require full-length PAD4

Since PAD4^LLD^ transgenic plants were as resistant as Col-0 against GPA, we tested if the PAD4^LLD^ domain also functions in basal and/or TNL-triggered pathogen immunity. For this, we measured TNL ETI using the biotrophic pathogen *Hyaloperonospora arabidopsidis* (*Hpa*) isolate EMWA1, which is recognized in Col-0 by the TNL *RPP4* (*RESISTANCE TO PERONOSPORA PARASITICA4*) [Van der Biezen et al., 2002; Asai et al., 2018]. Col-0, PAD4^WT^ and PAD4^S118A^ lines were fully resistant to *Hpa* EMWA1, as measured by conidiophore production (Figure 3A). By contrast, PAD4^LLD^ transgenic lines were fully susceptible with conidiophore production and macroscopic disease and microscopic *Hpa* colonization phenotypes resembling a *pad4-1/sag101-3* mutant (Figure 3A & Figure S3).

**Figure 3.**
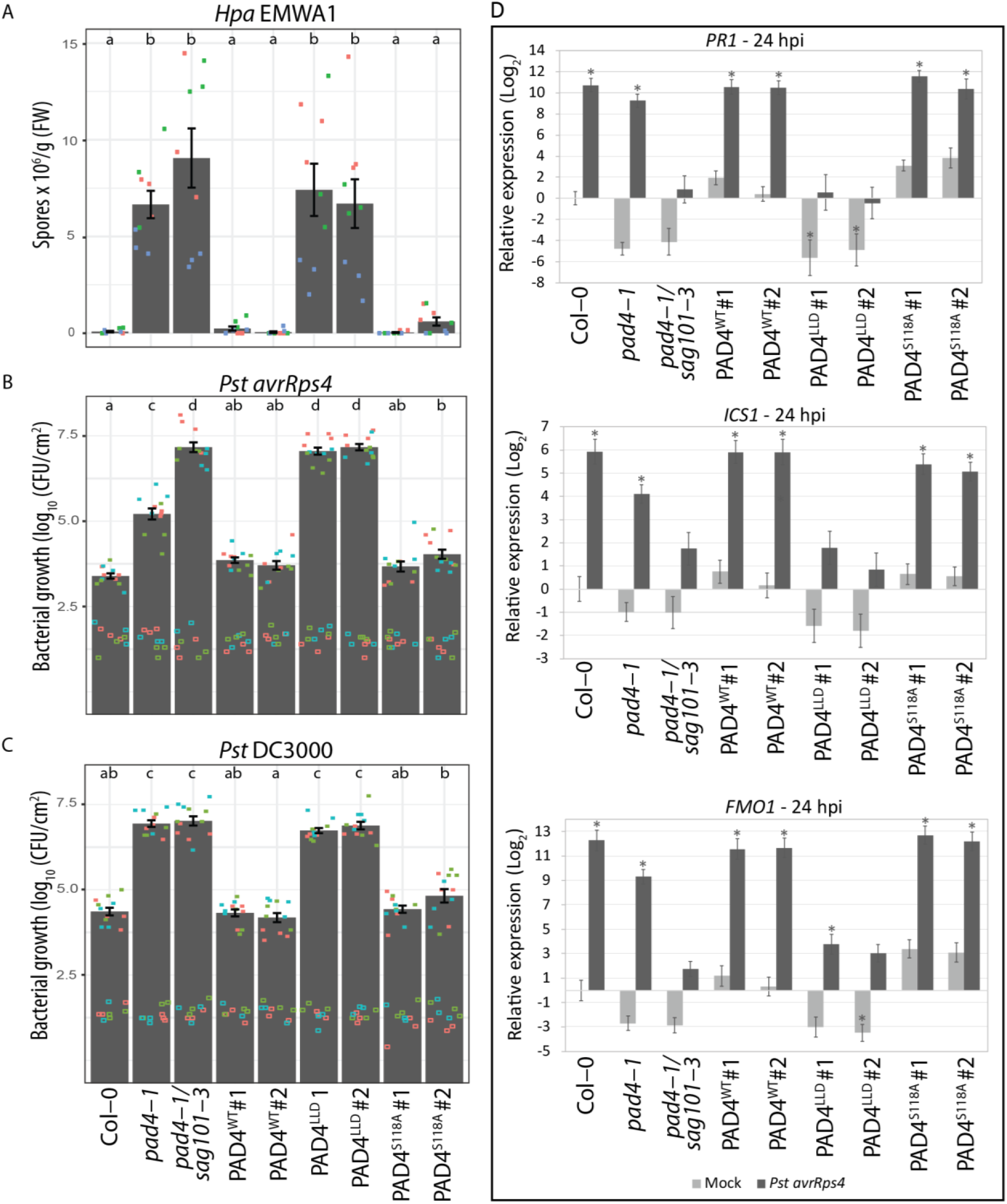
PAD4^LLD^ is not functional in Arabidopsis ETI and basal immunity. **A**. TNL (RPP4) ETI assay in Arabidopsis independent transgenic lines with wild-type and mutant controls, as indicated. *Hpa* EMWA1 conidiospores on leaves were quantified at 6 dpi in three independent experiments (squares; n=9). Col-0 (resistant), *pad4*-1 (susceptible) and *pad4-1/sag101-3* (susceptible) functioned as controls. Squares of the same color represent three biological replicates in an independent experiment. Bars represent mean of three experimental replicates ± SE. Differences between genotypes were determined using ANOVA (Tukey-HSD, *p* <0.01), letters indicate significance class. **B**. TNL (RRS1-S/RPS4) ETI assay in the same Arabidopsis independent transgenic and control lines as in A. Four-week old Arabidopsis plants were syringe infiltrated with *Pst avrRps4* (OD_600_ = 0.0005) and bacterial titers were determined at 0 dpi (empty squares; n=8-9) and 3 dpi (filled squares; n=11-12). Squares of the same color represent 2-3 (day 0) or 3-4 (day 3) biological replicates in an independent experiment. Bars represent mean of three experimental replicates ± SE. Differences between genotypes were determined using ANOVA (Tukey-HSD, *p* <0.01), letters indicate significance class. **C**. Infection assay was performed with basal immunity triggering *Pst* DC3000 (OD_600_ = 0.0005). Experimental set-up and statistical analysis as in B. **D**. Transcript abundance determined by RT-qPCR in 4-week old Arabidopsis plants syringe-infiltrated with either buffer (mock, grey bars) or *Pst avrRps4* (black bars) (24 hpi). Data are pooled from three independent experiments, with two to three biological replicates per experiment (n = 6–9). *PATHOGENESIS RELATED1* (*PR1*), *ISOCHORISMATE SYNTHASE1* (*ICS1*), and *FLAVIN MONOOXYGENASE1* (*FMO1*) transcript abundances were measured relative to *ACTIN2* (*ACT2*). Relative expression and significance level is set to Col-0 mock-treated samples. Differences between genotypes were determined using ANOVA (Tukey-HSD), asterisks indicate *p* < 0.01.

Further, we tested PAD4^LLD^ function in TNL (RRS1-S/RPS4) ETI to *Pst avrRps4* and in basal immunity to virulent *Pst* DC3000. In basal immunity, *pad4-1* is as susceptible as *pad4-1/sag101-*3 while in ETI, *pad4-1* displays intermediate susceptibility between Col-0 and *pad4-1/sag101-*3 (Figure 3B & 3C) [Feys et al., 2005; Wagner et al., 2013]. In line with published data, PAD4^S118A^ was as resistant as Col-0 and PAD4^WT^ in both basal immunity and ETI (Figure 3B & 3C), consistent with previous findings that the PAD4 S-D-H predicted catalytic triad is not required for pathogen immunity [Louis et al., 2012; Wagner et al., 2013]. PAD4^LLD^ lines were fully susceptible to *Pst* DC3000 and *Pst avrRps4*, with bacterial titers comparable to *pad4-1/sag101-3* (Figure 3B & 3C), indicating that PAD4^LLD^ is not able to confer basal immunity or ETI. Also, PAD4^LLD^ expressing plants and *pad4-1/sag101-3* failed to induce expression of defense marker genes 24 hpi with *Pst avrRps4*, indicating that PAD4^LLD^ is unable to signal in TNL ETI (Figure 3D-F & Figure S4). Taken together, the *Hpa* and *Pst* infection data show that PAD4^LLD^ is non-functional in pathogen basal immunity and ETI, in stark contrast to its resistance activity against GPA.

## Discussion

PAD4 controls Arabidopsis defenses against pathogens and aphids, playing major roles with EDS1 in basal and effector-triggered immunity, and an *EDS1*-independent role in resistance to GPA [Bhandari et al., 2019; Cui et al., 2017; Cui et al., 2018; Glazebrook et al., 1997; Lapin et al., 2019; Louis et al., 2012; Pegadaraju et al., 2007; Rietz et al., 2011; Wagner et al., 2013]. In this study, we investigated the contribution of the PAD4^LLD^ to these different defense outputs. Analysis of PAD4^LLD^ *in planta* shows that it accumulates to much lower levels than full-length PAD4 and has lost binding to EDS1. Strikingly, PAD4^LLD^ confers complete GPA resistance (Figure 2), but is non-functional in resistance to *Hpa* and *Pst* pathogens (Figure 3). Thus, PAD4 appears to rely solely on its LLD for controlling GPA infestation, whereas its LLD and EP domains are necessary for ETI and basal immunity against bacterial and oomycete pathogens. These data suggest there are domain specific signaling functions of Arabidopsis PAD4.

Recent studies suggest that the N-terminal LLDs of Arabidopsis EDS1 and PAD4 act as a scaffold, enabling the C-terminal EP domains to interact and orchestrate downstream immune signaling as a heterodimer [Bhandari et al., 2019; Lapin et al., 2019; Wagner et al., 2013]. By testing the PAD4^LLD^ without its EP domain, our data show that PAD4, like EDS1 [Bhandari et al., 2019], requires its EP domain for immunity signaling. By contrast, the PAD4^LLD^ is sufficient to limit GPA proliferation, thus highlighting a role of the PAD4^LLD^ and its α/β-hydrolase catalytic triad as a minimal functional unit in GPA resistance. *At*EDS1 and *At*PAD4 proteins mutually stabilize each other [Feys et al., 2001; Feys et al., 2005; Rietz et al., 2011; Wagner et al., 2013]. The fact that interaction between PAD4^LLD^ and EDS1 is greatly diminished compared to interaction between full-length PAD4 and EDS1 (Figure 1D), tallies with the observation that *PAD4*-dependent GPA resistance is independent of *EDS1* [Pegadaraju et al., 2007].

The PAD4^LLD^ adopts an α/β hydrolase fold with a core S-D-H predicted catalytic triad. The α/β hydrolase family catalyzes a variety of enzymatic reactions such as esterification, hydrolysis and acyl transfer [Rauwerdink & Kazlauskas, 2015]. The S-D-H predicted catalytic triad of PAD4 is dispensable for immune signaling against *Hpa* and *Pst*, but required for GPA resistance [Louis et al., 2012; Wagner et al., 2013]. In the *At*PAD4 structural model, this triad of residues is solvent-accessible (Figure 1A & 1B), suggesting a plausible catalytic function. However, this applies only to Brassicaceae PAD4 proteins, as beyond the Brassicaceae clade PAD4 contains an insertion, which forms a “lid” covering the S-D-H triad similar to that in *At*EDS1, rendering it inaccessible to the solvent [Wagner et al., 2013]. Such helical loop structures extending from the β-sheet scaffold have been found to regulate the enzymatic activity of inactive-state triacylglycerol lipases [Khan et al., 2017]. Hence, it is possible that the PAD4 S-D-H triad functions differently outside the Brassicaceae clade [Wagner et al., 2013]. Critically, all three residues in the catalytic triad are required for hydrolase activity [Rauwerdink & Kazlauskas, 2015]. Since loss of H^229^ does not affect *At*PAD4-mediated deterrence of GPA [Louis et al., 2012], it is more likely that PAD4 involvement in Arabidopsis defense against the GPA does not rely on a canonical hydrolase activity.

Alternatively, the S-D-H triad in PAD4 could function as a receptor ligand-binding domain, a common feature of α/β hydrolase fold proteins [Mindrebo et al., 2016]. For example, the Arabidopsis karrikin receptor *At*KAI2 (KARRIKIN INSENSITIVE 2) uses its catalytically inactive S-D-H triad for ligand recognition [Guo et al., 2013]. Catalytically inactive rice (*Os*) and *At*GID1 (GIBBERELLIN (GA) INSENSITIVE DWARF1) uses a modified triad (S-D-V) to bind bioactive GA molecules, indicating that the histidine, which is required for catalytic activity, can be replaced by another residue for functional ligand binding [Murase et al., 2008; Rauwerdink & Kazlauskas, 2015; Shimada et al., 2008]. Upon binding to GA, a conformational change in *At*GID1 results in the assembly of a SCF^GID1^ (SKP-Cullin-F-box^GID1^) complex and ubiquitination of DELLA proteins marking them for proteasome-mediated degradation [Murase et al., 2008]. Together with the data presented here, these examples highlight the possibility that the PAD4^LLD^ domain serves as a ligand-binding surface in a protein signaling complex, rather than a lipase.

The inactivity of PAD4^LLD^ in basal and effector-triggered immunity is unlikely to be attributed to PAD4^LLD^ instability, as it is sufficient for resistance against GPA, unless the PAD4^LLD^ fails to reach sufficient amounts needed for pathogen resistance in certain cells or tissues. Very low levels of protein were sufficient for EDS1 function in pathogen immunity [Bhandari et al., 2019; Suttmann et al., 2016; Wagner et al., 2013], and we presume this is also the case for PAD4, since EDS1 and PAD4 are functional as a heterodimer. A more plausible explanation for the susceptibility of PAD4^LLD^ might be (i) its inability to form a heterodimer with EDS1 (Figure 1A & 1D; Wagner et al., 2013) and (ii) the lack of an EP domain. Both are essential in EDS1 for immunity to *Pst* and *Hpa* infection and for rapid transcriptional up-regulation of defense genes in ETI against *Pst avrRps4* [Bhandari et al., 2019; Lapin et al., 2019; Wagner et al., 2013]. The EDS1 EP domain interface lining a cavity formed with PAD4 in the heterodimer is necessary for Arabidopsis EDS1 signaling [Bhandari et al., 2019; Lapin et al., 2019]. An aligned EP domain α-helix was identified in the EDS1 heterodimer partner, SAG101, as being essential for eliciting host cell death in TNL ETI responses [Gantner et al., 2019; Lapin et al., 2019]. This also might be true for PAD4, because mutations at an EDS1-like surface in PAD4 lying outside the cavity did not compromise immunity [Bhandari et al., 2019]. Future studies will test whether the PAD4 EP domain surface lining the heterodimer cavity is also crucial for EDS1-PAD4 pathogen immunity.

Our analysis of the LLD of Arabidopsis PAD4 demonstrates a domain-specific partitioning of defense functions - with the PAD4^LLD^ being necessary and sufficient for limiting GPA infestation, and the EP domain (with the LLD) mediating immunity signaling against *Pst* and *Hpa* (Figure 4). While the two PAD4 domains clearly have distinct roles, instability and inactivity of the PAD4 EP domain without the LLD makes it difficult to assess whether PAD4 is a bipartite immune regulator or moonlighting in GPA resistance. This study of the PAD4^LLD^ paves way for molecular dissection of the diverse roles of PAD4 in biotic stress resistance.

**Figure 4.**
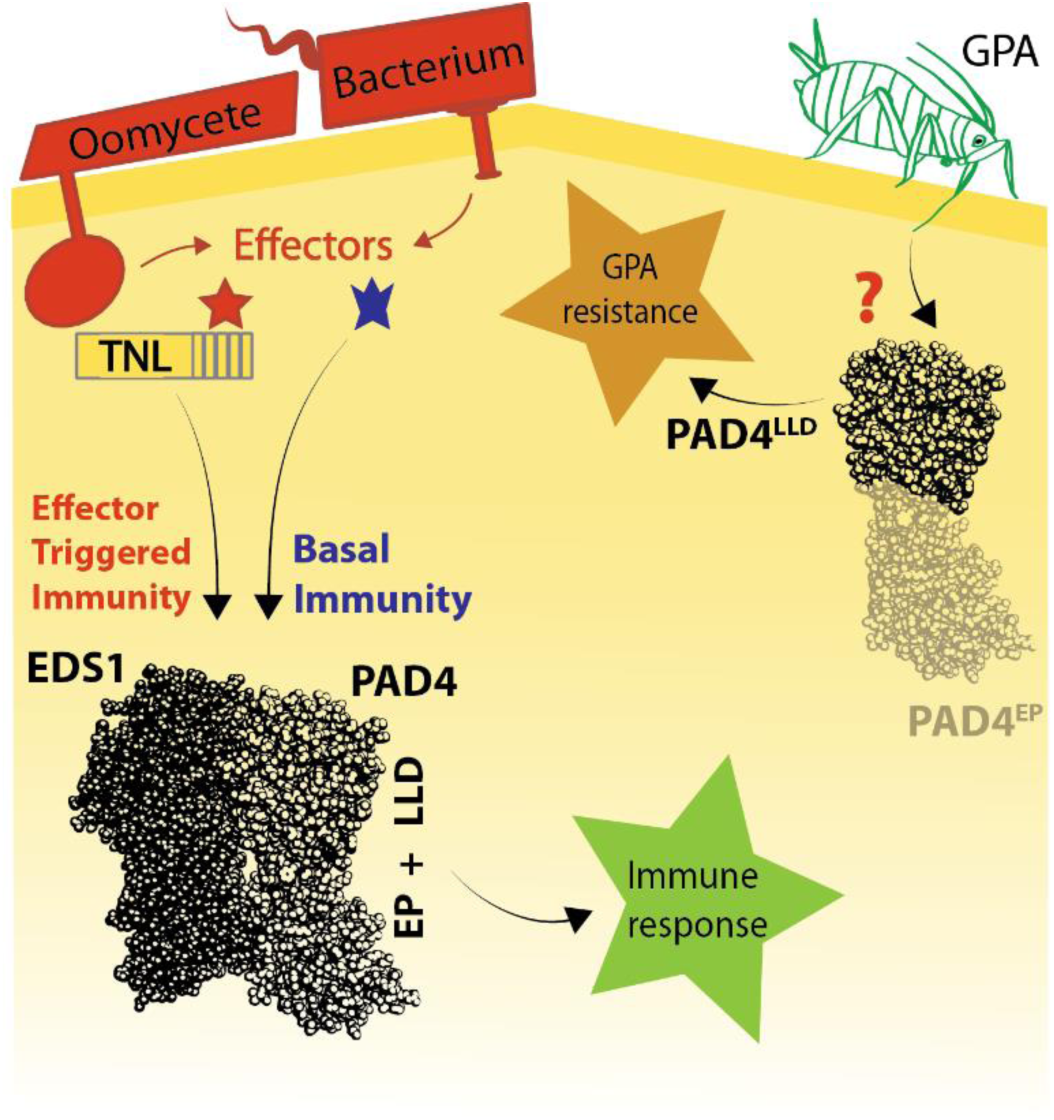
Domain-specific roles of Arabidopsis PAD4 in immunity. Schematic showing separable activities of PAD4^LLD^ and PAD4^LLD^ + EP domain. Upon infection by bacteria or oomycetes, the EDS1-PAD4 heterodimer is activated via TNLs in ETI or by other signals in basal immunity, leading to a pathogen immune response. PAD4 requires both the LLD and EP domains to function in basal immunity and ETI. In resistance to GPA, PAD4 is activated through an unknown but *EDS1*-independent mechanism that restricts aphid infestation. PAD4^LLD^ is sufficient to limit GPA independently of interaction with EDS1.

## Author contributions

JAD, DDB, JEP designed the study; JAD, LuD, MP and LA performed experiments, JAD and JEP wrote the manuscript with inputs from DDB, LaD and JS; JEP, LaD and JS provided funding.

## Acknowledgments

We would like to thank Karsten Niefind for structural insights and Neysan Donnelly for help editing the manuscript.

## Materials and Methods

### Plant materials, growth conditions and pathogen strains

Arabidopsis *pad4-1, sag101-3*, and *eds1-2* mutants are in the Col-0 background and were previously described, as were *pEDS1:EDS1-SDH*::*pPAD4:PAD4-S118A* (in *eds1-2/pad4-1*) and *pPAD4:StrepII-YFP-PAD4* (in *pad4-1/sag101-3*) transgenic lines (See table S1 for primers) [Bhandari et al., 2019; Wagner et al., 2013]. *Pseudomonas syringae* pv. *tomato* (*Pst*) strain DC3000 and *Pst avrRps4* were described previously (Cui et al., 2017). Plants were grown on soil in a controlled environment and insect-free chambers under a 10 h light/14 h dark regime (PAR: 100–150 μmol/m^2^/s) at 22 °C and 60% relative humidity.

### Pathogen infection assays

For bacterial growth assays, *Pst avrRps4* (OD_600_=0.0005) in 10 mM MgCl_2_ was hand-infiltrated into leaves of 4-week-old plants. Bacterial titers were measured at 3 h post-infiltration (day 0) and 3 d, as described previously (Feys et al., 2005). Each biological replicate consisted of three leaf disks from different plants and data shown in each experiment are compiled from three to four biological replicates. Statistical analysis was performed using one-way ANOVA with multiple testing correction using Tukey’s HSD (*p*-value as described in figure legend).

For gene expression analysis, leaves of 4-week-old plants were hand-infiltrated with mock (10 mM MgCl_2_) or bacteria (OD_600_= 0.005) and samples were taken at 24 hpi. *ACT2* was used as a reference gene (See Table S1 for primers). Data shown are results from three independent experiments each with two to three biological replicates. For protein accumulation assays, leaves from 4-week-old plants were hand-infiltrated with buffer (mock, 10 mM MgCl_2_) or bacteria (OD_600_= 0.005) and samples harvested by pooling leaves from at least three different plants.

*Hpa* isolate EMWA1 was sprayed onto 2.5 week-old plants at 4 × 10^4^ spores/ml dH_2_O. *Hpa* infection structures and plant host cell death were visualized using lactophenol trypan blue staining (Muskett et al., 2002) and imaged by light microscopy (Zeiss Axio Imager). To quantify *Hpa* sporulation on leaves, three pots with ∼ 10 plants per genotype were infected and treated as a biological replicate. Plants were harvested at 6 dpi, fresh weight was determined. Conidiospores were suspended in 5 ml dH_2_O and counted under a light microscope using a Neubauer counting chamber.

### Aphid no-choice bioassay

For each biological replicate five one-day-old nymphs were released onto the center of a 17-day-old plant. The total number of aphids (adult + nymphs) per biological replicate were counted 11 days post infestation. Each independent experimental replicate consisted of 10 biological replicates per genotype [Nalam et al., 2018].

### Plasmid constructs

The pENTR/D-TOPO *PAD4* vector used for site-directed mutagenesis was cloned from cDNA and is described [Wagner et al., 2013]. PAD4^LLD^ was obtained by site-directed mutagenesis on pENTR/D-TOPO *PAD4* according to the QuikChangeII site-directed mutagenesis manual (Agilent) (See Table S1 for primers). Mutated *PAD4* and *EDS1* entry clones [Bhandari et al., 2019; Wagner et al., 2013] were verified by sequencing and recombined by an LR reaction into a pAM-PAT-based binary vector backbone [Witte et al., 2004]. Split-luciferase lines were created by LR reaction between gateway-compatible split-luciferase binary vectors [Gehl et al., 2011] and *PAD4* and *EDS1* entry clones [Bhandari et al., 2019; Wagner et al., 2013].

### Generation of transgenic Arabidopsis plants

Stable transgenic lines were generated by transforming a binary expression vector (containing Basta resistance) into Arabidopsis null mutant *pad4-1/sag101*-3 [Wagner et al., 2013], using *Agrobacterium*-mediated floral dipping (*Agrobacterium tumefaciens* GV3101 PMP90 RK) [Clough and Bent, 1998]. After selecting single-insert, homozygous transgenic lines, all lines were genotyped by sequencing for the presence of the correct PAD4 transgene (PAD4^WT^, PAD4^LLD^ or PAD4^S118A^) before performing pathogen assays.

### Transient expression in *N. benthamiana*

Transient expression in *N. benthamiana* was performed by co-infiltrating *Agrobacterium* cells carrying constructs at an OD_600_ of 0.4–0.6 in a 1:1 ratio. Before syringe infiltration, *A. tumefaciens* cells were incubated for 3h at 28°C in induction buffer (150 µM acetosyringone,10 mM MES pH5.6, 10 mM MgCl_2_) and shaked at 650 rpm in an Eppendorf Thermomixer. *N. benthamiana* leaf samples were harvested at 3 dpi and snap frozen in liquid nitrogen and stored at −80°C.

### Protein extraction, immunoprecipitation (IP) and Western blotting

Total leaf extracts were processed in extraction buffer (50 mM Tris pH7.5, 150 mM NaCl, 10 % (v/v) glycerol, 2 mM EDTA, 5 mM DTT, 0.1 % Triton X-100 and protease inhibitor (Roche, 1 tablet per 50 ml)). Lysates were centrifuged for 20 min, 21,000 x *g* at 4 °C. Supernatant was used as input sample (50 μl). Immunoprecipitations were conducted by incubating the input sample (1.2 mL) with 10 μl GFP TrapMA beads (Chromotek) for 3 h at 4 °C. Beads were collected using a magnetic rack and washed four times in extraction buffer. Protein or IP samples were boiled at 96 °C in 2x Laemmli buffer for 10 min. Proteins were separated by SDS-PAGE and analyzed by immunoblotting using α-GFP (Sigma Aldrich, 11814460001) or α-FLAG (Sigma Aldrich, F7425) primary antibodies and secondary antibodies coupled to Horseradish Peroxidase (HRP, Sigma Aldrich) for protein detection on blots.

### Luciferase Assay

All tested co-expression constructs were transiently expressed on one leaf. Three leaf disks (0.4 cm diameter) from three independent leaves were pooled per biological replicate and processed in reporter lysis buffer (Promega; E1500, + 150 mM Tris, pH 7.5). Samples were mixed in a 1:1 ratio with substrate (Promega; E1531) and luminescence was measured. Absolute luminescence, *i*.*e*. absolute luciferase activity was used as a proxy for protein-protein interaction intensity.

## Supplemental Material

**Figure S1.**
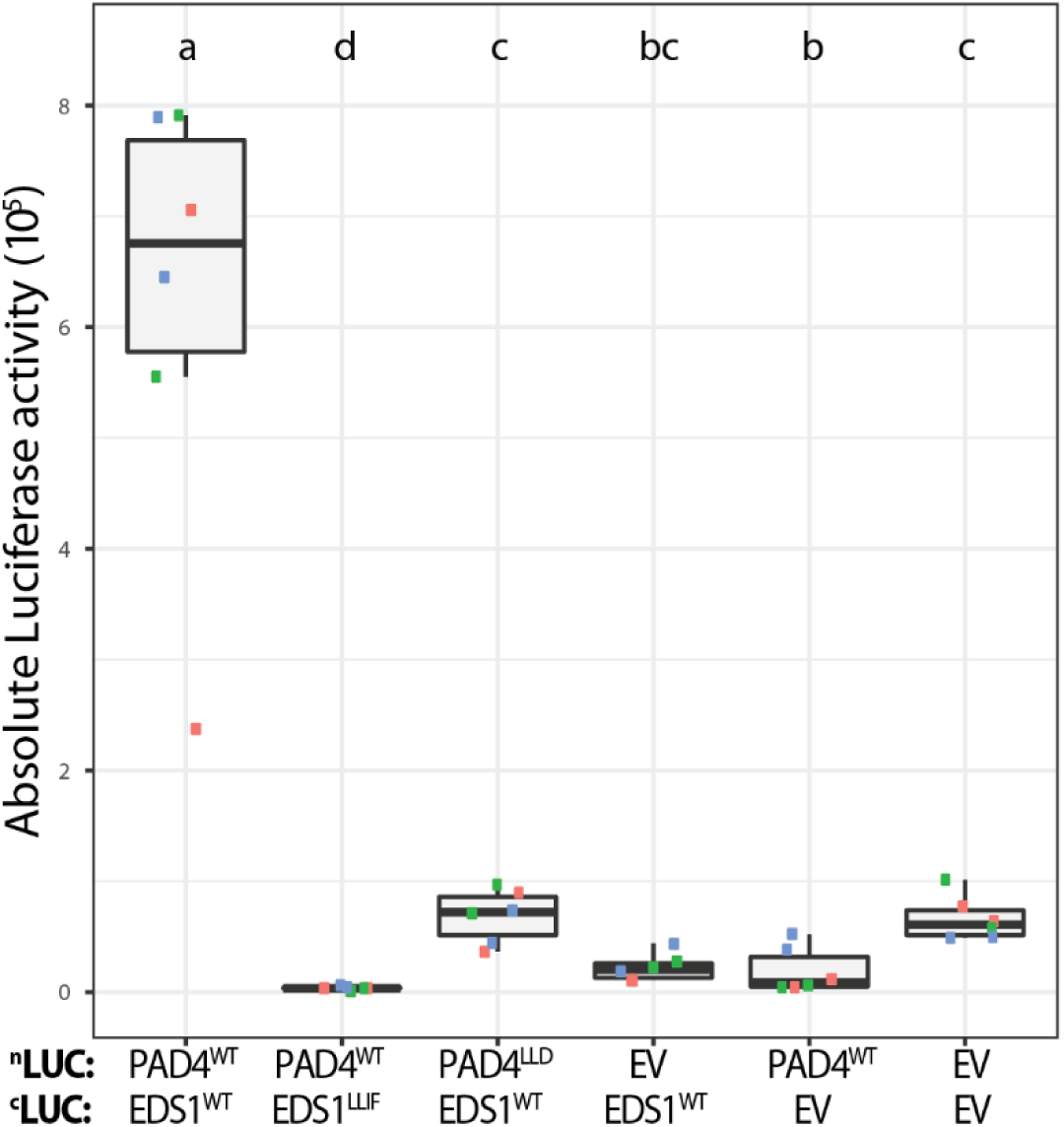
Split-luciferase interaction assay. Absolute luciferase (LUC) activity from transiently co-expressed N-LUC or C-LUC constructs (35S promoter) in *N. benthamiana*. Data are pooled from three independent experiments with two biological replicates per experiment (n = 6). Error bars = SEM. Letters indicate statistical significance as determined by one-way ANOVA with multiple testing correction using Tukey-HSD; *p* < 0.01.

**Figure S2.**
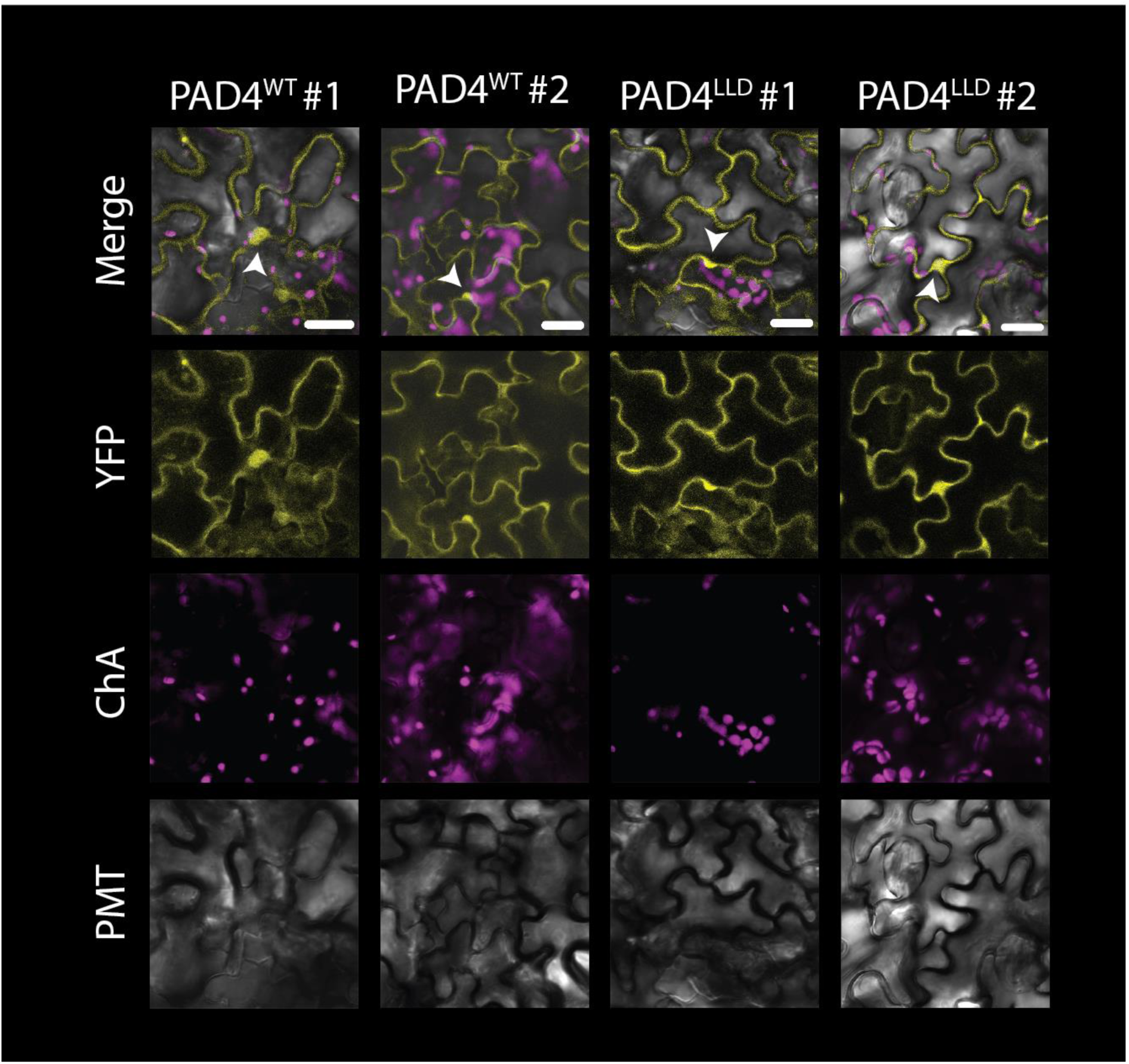
PAD4 localization at 24 hpi with *Pst* AvrRps4. Nucleocytoplasmic localization of YFP-PAD4^WT^ and YFP-PAD4^LLD^ in Arabidopsis transgenic lines (24 hpi, *Pst avrRps4*). To determine PAD4^LLD^ localizations, confocal microscope sensitivity was enhanced to enable its detection. White arrowheads indicate nuclei and white bars correspond to 20 µm. Similar results were obtained in two independent replicates in two biological replicates (n=4).

**Figure S3.**
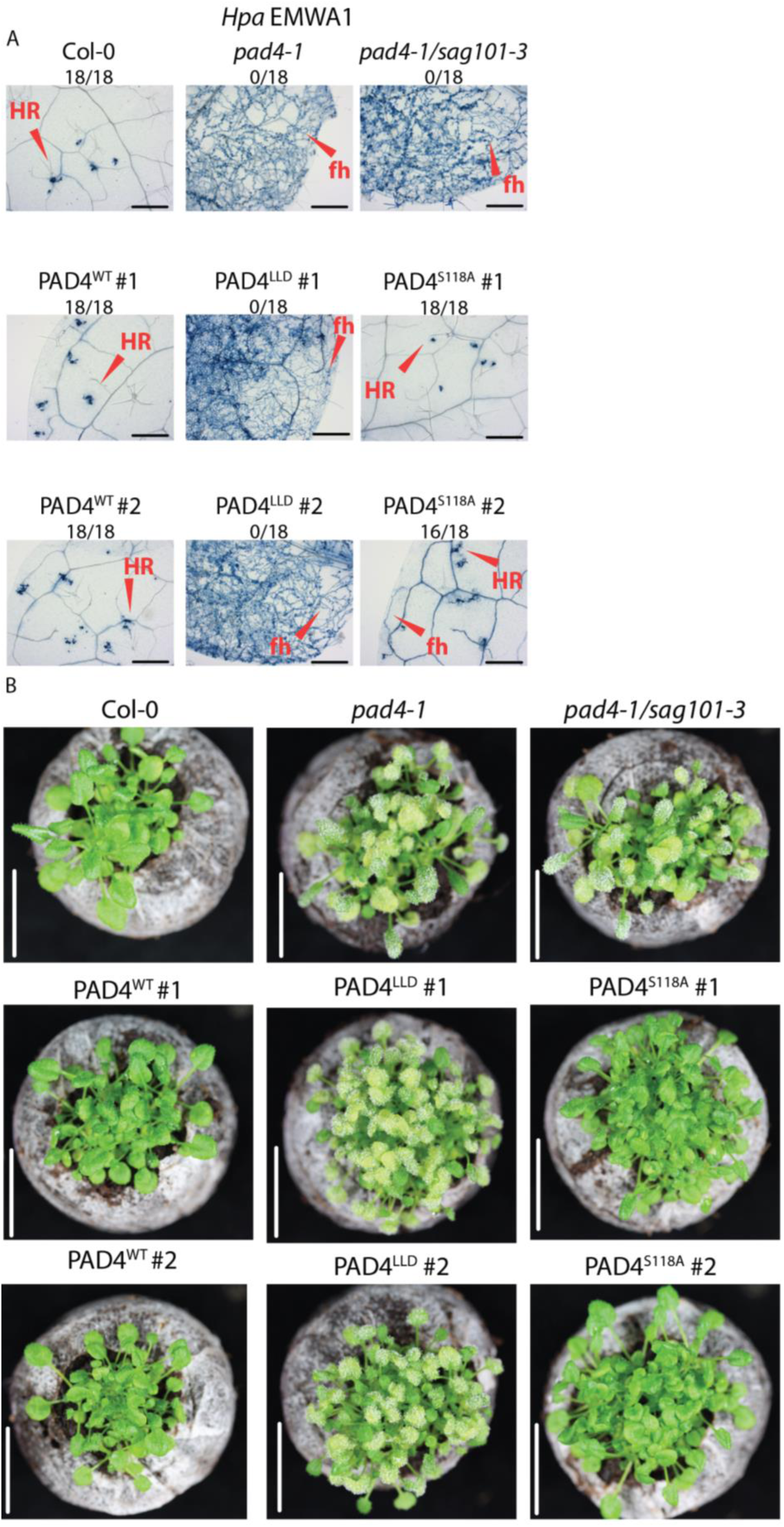
Microscopic and macroscopic disease phenotypes of *Hpa* EMWA1-infected Arabidopsis plants. **A**. Microscopic immunity phenotypes of 3-week-old Arabidopsis lines, as indicated, at 6 dpi with *Hpa* isolate EMWA1 (recognized by TNL RPP4). Trypan blue-stained leaves showing free hyphae (fh) and hypersensitive cell death (Hypersensitive Response (HR)). Black bars represent 500 μm. Fractions (*e*.*g*. 18/18) indicate numbers of resistant leaves/total plants tested. Pictures are representative from three independent experimental replicates, > 6 leaves per replicate and > 30 infection sites per genotype. **B**. *Hpa* EMWA1-inoculated plants of the same lines as in A. Resistant plants look healthy at 6 dpi, whereas susceptible plants produce conidiospores and leaf chlorosis. White bars correspond to 2 cm.

**Figure S4.**
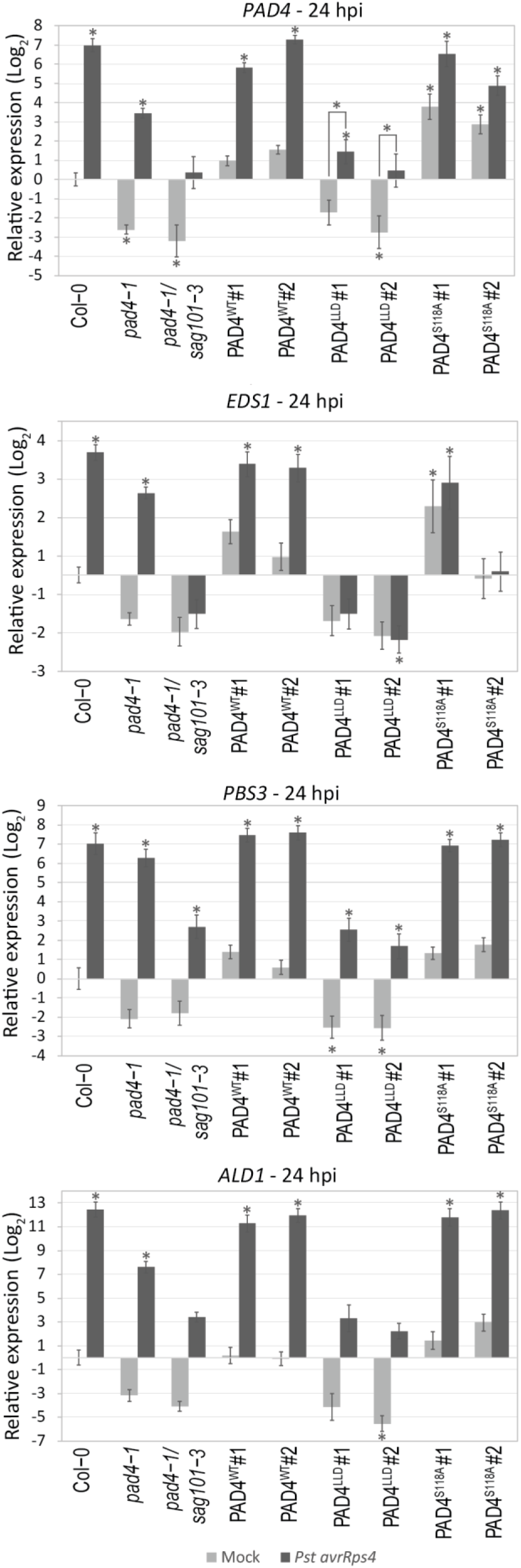
Relative expression of *EDS1*-dependent defense marker genes. Transcript abundance was determined by RT-qPCR in 4-week old Arabidopsis plants of the indicated lines syringe infiltrated with buffer-(mock, grey bars) or *Pst avrRps4* (black bars) (24 hpi). Data are pooled from three independent experiments with two to three biological replicates per experiment (n = 6–9). Transcript abundances of *PAD4, EDS1, AVRPPHB SUSCEPTIBLE 3* (*PBS3*) and *GD2-LIKE DEFENSE RESPONSE PROTEIN 1* (*ALD1*) were measured relative to *ACT2*. Relative expression and significance level are set to Col-0 mock-treated samples. Bars represent mean of three experimental replicates ± SE. Differences between genotypes were determined by using ANOVA (Tukey-HSD, *p* <0.01), letters indicate significance class and asterisk indicates *p* < 0.01.

**Table S1.**
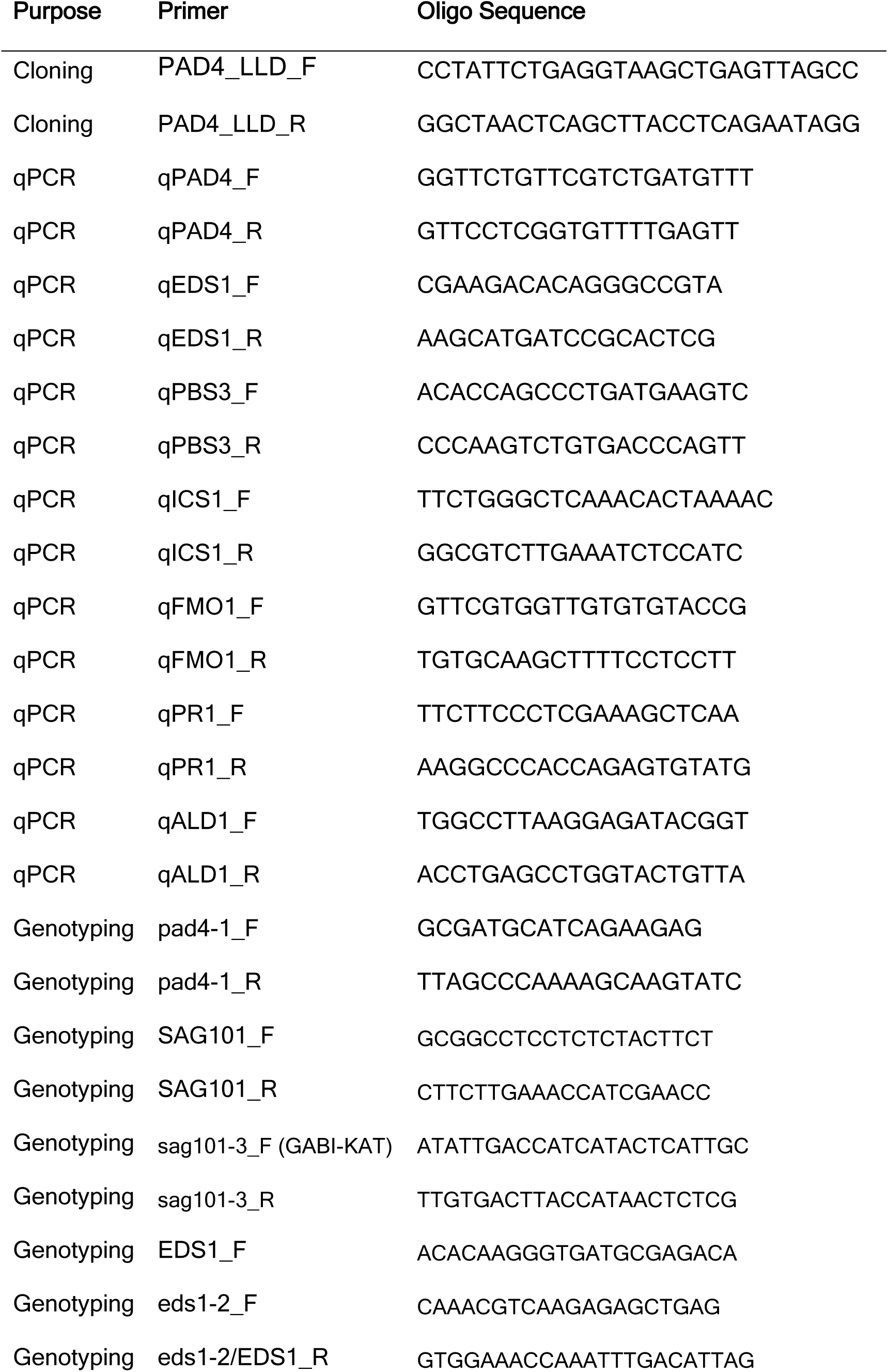
List of DNA primers used in this study.

## Notes

Funding: This work was supported by the Max Planck Society and Deutsche Forschungsgemeinschaft (DFG) Grants: DFG-ANR Trilateral ‘RADAR’ grant (JAD, LaD (ANR-15-CE20-0016-01), JEP), CRC670 (DDB, JEP), and MP was supported by a scholarship from the University of North Texas.

